# Mechanical transmission of African swine fever virus by *Stomoxys calcitrans*: insights from a mechanistic model

**DOI:** 10.1101/2020.07.10.179853

**Authors:** Vergne Timothée, Andraud Mathieu, Bonnet Sarah, De Regge Nick, Desquesnes Marc, Fite Johanna, Etore Florence, Garigliany Mutien-Marie, Jori Ferran, Lempereur Laetitia, Le Potier Marie-Frédérique, Quillery Elsa, Saegerman Claude, Vial Laurence, Bouhsira Emilie

## Abstract

African swine fever (ASF) represents a global threat with huge economic consequences for the swine industry. Even though direct contact is likely to be the main transmission route from infected to susceptible hosts, recent epidemiological investigations have raised questions regarding the role of hematophagous arthropods, in particular the stable fly (*Stomoxys calcitrans*). In this study, we developed a mechanistic vector-borne transmission model for ASF virus (ASFV) within an outdoor domestic pig farm in order to assess the relative contribution of stable flies to the spread of the virus. The model was fitted to the ecology of the vector, its blood-feeding behaviour and pig-to-pig transmission dynamic. Model outputs suggested that in a context of low abundance (<5 flies per pig), stable flies would play a minor role in the spread of ASFV, as they are expected to be responsible for around 10% of transmission events. However, with abundances of 20 and 50 stable flies per pig, the vector-borne transmission would likely be responsible for almost 30% and 50% of transmission events, respectively. In these situations, time to reach a pig mortality of 10% would be reduced by around 26% and 40%, respectively. The sensitivity analysis emphasised that the expected relative contribution of stable flies was strongly dependent on the volume of blood they regurgitated and the infectious dose for pigs. This study identified crucial knowledge gaps that need to be filled in order to assess more precisely the potential contribution of stable flies to the spread of ASFV, including a quantitative description of the populations of hematophagous arthropods that could be found in pig farms, a better understanding of blood-feeding behaviours of stable flies, and the quantification of the probability that stable flies partially fed with infectious blood transmits the virus to a susceptible pig during a subsequent blood-feeding attempt.

## INTRODUCTION

African swine fever (ASF) represents a global threat with huge economic consequences for the swine industry (Sánchez-Cordón et al., 2018; Beltran-Alcrudo et al., 2019). The large-scale spread through Eastern Europe since 2007 reinforces the need to understand the epidemiological determinants of virus transmission (Cwynar et al., 2019). Direct contact is likely to be the main transmission route from infected to susceptible hosts (Guinat, Gogin et al., 2016; Halasa et al., 2016), although indirect short-distance transmission was also highlighted as being non-negligible, especially in areas with high density of small free-range pig herds (Andraud et al., 2019). Human activity is certainly one of the most important drivers of virus introduction into pig farms, due to commercial exchanges of live animals and sharing of material between neighbouring farms (Lichoti et al., 2016; Kukielka et al., 2017; Beltran-Alcrudo et al., 2019). In African regions where ASF is endemic, soft ticks of the species *Ornithodoros moubata* were identified as competent vectors for ASF virus (ASFV), contributing to the sylvatic cycle of ASF in warthogs (Quembo et al., 2016, 2018; Penrith et al., 2019). *Ornithodoros moubata* ticks are nevertheless almost absent from Palearctic ecozones and cannot be considered as playing a role in the spread of the virus in Europe. Soft ticks of the species *O. erraticus* are present in the Iberian peninsula where ASF was endemic for three decades (Pérez-Sánchez et al., 1994; Vial et al., 2018), but experimental infections have suggested that they might be much less efficient in transmitting the Georgia 2007/01 ASFV strain to pigs than *O. moubata* (Oliveira et al., 2019). Epidemiological data from Eastern Europe highlighted concomitant outbreaks in wild boars and domestic pigs, showing a viral spread at the interface between these two host types (Podgórski and Śmietanka, 2018; Cwynar et al., 2019). Direct contacts between animals was clearly an important determinant, especially for free-range herds. Hematophagous arthropods, that can potentially bite different hosts to complete a blood meal, could also play a role in this between-species transmission, particularly in the introduction of the virus into high-biosecurity farms (Herm et al., 2019; Fila and Woźniakowski, 2020). A strong seasonality of ASF outbreaks in domestic pigs with a peak in summer was observed in Estonia, Poland, Latvia and Lithuania raising questions about the potential role of hematophagous insects presenting a similar seasonal activity (EFSA, 2020). For these reasons, the European Food Safety Authority has strongly emphasized the need to improve our understanding of the potential role of blood-feeding arthropods, other than soft ticks, in the spread of ASFV (Miteva et al., 2020).

In the early 1980s, experimental transmissions have demonstrated that stable flies (*Stomoxys calcitrans*) could transmit ASFV to susceptible pigs one to 24 hours after an infective blood meal (Mellor et al., 1987), and therefore could act as mechanical vectors of ASFV. Recently, infectious virus was isolated from the body of stable flies three to 12 hours after *in vitro* infections (Olesen, Hansen et al., 2018), and experimental infection was demonstrated after ingestion of contaminated stable flies (Olesen, Lohse et al., 2018). In addition, several observations suggest that stable flies can be present both in pig farms and in forest areas where wild boar and domestic pig farms can be in close proximity (Fischer et al., 2001; Petrasiunas et al., 2018). Following these empirical findings, a recent prioritization study based on expert-opinion elicitation ranked stable flies as the most probable blood-feeding arthropod to be a mechanical vector of ASFV in metropolitan France (Saegerman et al., 2020). This rank was justified by the fact that stable flies often need to feed on different hosts to complete a blood meal, that they can complete blood meals very regularly within a day (Kunz and Monty, 1976), and that they have been shown to be able to regurgitate blood stored in their crop during blood feeding attempts (Coronado et al., 2004; Baldacchino et al., 2013).

Estimating the relative contribution of blood-feeding arthropods in the transmission of ASFV is crucial for developing efficient biosecurity measures, including vector control. Mechanistic models may help to address this issue through the evaluation of the relative impact of this potential transmission route both in terms of virus introduction probability and within-farm spread (Turner et al., 2012; Sumner et al., 2017).

The objectives of the study were to 1) develop a mechanistic modelling framework for vector-borne transmission of the Georgia 2007/01 ASFV strain within a pig farm, 2) assess the relative contribution of mechanical transmission by stable flies in the spread of ASFV in an outdoor pig farm and 3) identify the critical knowledge gaps to drive future research.

## MATERIALS AND METHODS

The potential role of stable flies in the mechanical transmission of ASFV was investigated in the context of an outdoor domestic pig farm, in which stable flies can complete their full development cycle since they usually have direct access to pig manure. The pig farm was assumed to host 200 pigs, what is representative of outdoor domestic pig farms in France. Information related to the density of stable flies in pig farms is very scarce. To the best of the authors’ knowledge, (Moon et al., 1987) is the only published article that mentions stable flies abundance in pig facilities. They reported between three and seven stable flies per pig in confined nurseries. As a consequence, different scenarii of level of infestation in farms were considered in the model to cover low infestation levels (5 flies per pig) to high infestation levels (100 per pig, as easily observed on horses or cattle).

### Model formulation

To address the study’s objectives, we developed a mechanistic model of ASFV transmission that incorporated a vector compartment (Figure 1). Briefly, pigs started as susceptible individuals (S_p_, not yet infected). Upon infection, they entered the exposed compartment (E_p_, infected but not yet infectious) in which they stayed during a period averaging the latent period duration (μ). Subsequently, pigs stayed in the infectious compartment (I_p_) during a period averaging the infectious period duration (σ). Since the lethality rate of the Georgia 2007/01 ASFV strain is close to 100% (Dixon et al., 2020), we assumed that pigs died at the end of the infectious period. Stable flies started also as susceptible individuals (S_v_). Since we considered mechanical transmission only, we assumed that, upon infection, vectors entered directly into the infective compartment (I_v_) in which they stayed during a period averaging the infective period duration (τ). At the end of the infective period, they moved back to the susceptible state. The vector population size was assumed constant with a birth rate (δ) equalling the mortality rate. Pigs and stable flies were assumed to mix homogeneously.

**Figure 1:**
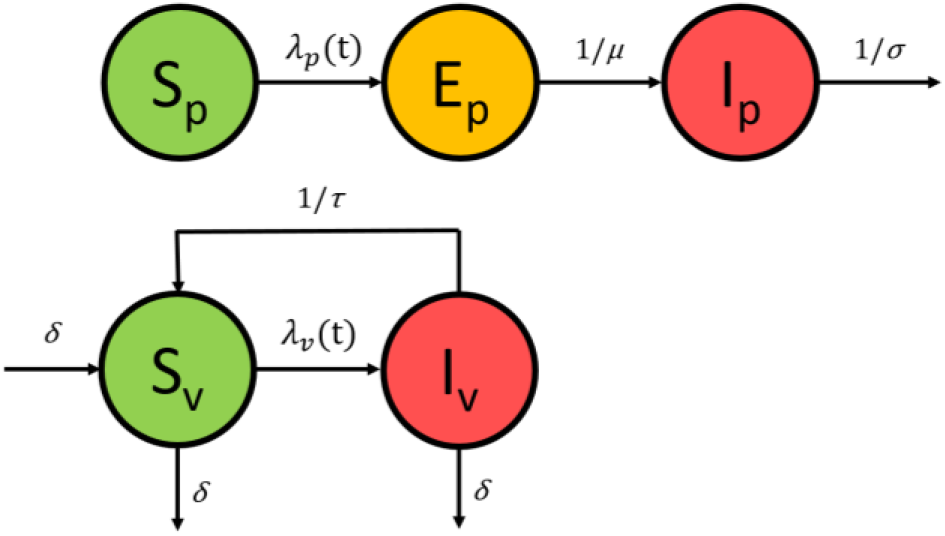
Schematic representation of the population-based model structure. Pigs and stable flies start as susceptible individuals (compartment S_p_ and S_v_, respectively) and become infected at rates **λ_p_(t)** and **λ_v_(t)**, respectively. Upon infection, pigs enter the exposed compartment (E_p_) in which they stay during a period averaging the incubation period duration (μ). Subsequently, they move to the infectious compartment (I_p_) where they stay until they die, during a period averaging the infectious period duration (σ). Infective stable flies remain as such until the end of their infective period duration (τ). At the end of the infective period, they moved back to the susceptible state. The stable fly population size was assumed constant with a birth rate (δ) equalling the mortality rate.

The force of infection exerted on a susceptible pig was given by

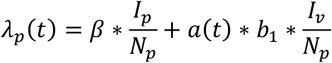

while the force of infection exerted on a susceptible vector was given by

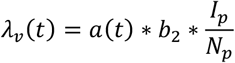

with *β* being the transmission rate between pigs, a(t) being the biting rate of stable flies as a function of time, b_1_ being the probability that a pig becomes infected if bitten by an infective vector, b_2_ being the probability that a vector becomes contaminated if it bites an infectious pig and N_p_ being the total number of live pigs present on site.

### Model parametrisation

Parameter values and associated references are provided in Table 1.

**Table 1:** Parameter values related to the model of ASFV mechanical transmission by S. calcitrans

| Parameter | Description | Value | Reference |
| --- | --- | --- | --- |
| $N_p$ | Initial herd size | 200 | Model scenario |
| $Ratio_{SP}$ | Number of stable flies per pig | 5, 10, 20, 50 and 100 | Model scenarii |
| $\beta$ | Daily transmission rate between pigs | Pert(0.2; 0.4; 0.6) day<sup>-1</sup> | Adapted from (Guinat, Gubbins et al., 2016) |
| $\mu$ | Average duration of the latent period in pigs | Pert(3; 4; 5) days | (Guinat, Gubbins et al., 2016) |
| $\sigma$ | Average duration of the infectious period in pigs | Pert(3; 7; 14) days | (Guinat, Gubbins et al., 2016) |
| $1/\delta$ | Average length of stay of stable flies in pig farms | 21 days | (Gilles, 2005) |
| $a(t)$ | Average biting rate by stable flies | Pert(0.4; 0.5; 0.6) hour<sup>-1</sup> if 8am < t < 8pm; 0 otherwise | See text for justification |
| $b_1$ | Probability that a pig becomes infected if bitten by an infective vector | $4.10^{-4}$ | See text for justification |
| $b_2$ | Probability that a vector becomes contaminated if it bites an infectious pig | 1 | See text for justification |
| $\tau$ | Average duration of the infective period in stable flies | 24 hours | See text for justification |

During a transmission experiment in a confined environment (Guinat, Gubbins et al., 2016), the daily transmission rate between pigs of the same pen was estimated at 0.6 (95% confidence interval: 0.3, 0.9) while the daily transmission rate between pigs of adjacent pens was estimated at 0.4 (95%CI: 0.1, 0.7). To account for the fact that this model was applied to an outdoor domestic pig farm in which contact rates were likely to be smaller than in the transmission experiment environment described in (Guinat, Gubbins et al., 2016), and that the layout of the farm was not considered, the daily pig-to-pig transmission rate was adjusted and assumed to be distributed according to a Pert distribution of parameters minimum=0.2, mode=0.4 and maximum=0.6.

Adult stomoxes live between two and four weeks on average (Gilles, 2005; Salem et al., 2012) and are able to remain in the same suitable environment their whole life if feeding and development criteria are met (represented here by access to pigs for feeding and to pig manure for lifecycle completion). Therefore, we assumed that stable flies could live on the outdoor domestic pig farm (and take blood meals) for an average duration of 21 days.

Because stable flies are actively looking for a blood meal during daytime (Berry and Campbell, 1985), it was considered that their biting activity period occurred between 8am and 8pm. Even though their activity can show some hourly variations during a day, this blood feeding pattern is not consistent across observations as it is highly dependent on the temperature (Baldacchino et al., 2013). Consequently, in the absence of more consistent evidence, the biting rate was assumed constant during the biting activity period. Stable flies feed frequently, with a time duration between two blood meals reported between four hours and four days (Salem et al., 2012). Due to the pain generated by a bite, stable flies’ blood meals are generally interrupted and split into five to 20 attempts, depending on the host reactivity. Therefore, it was assumed that stable flies could make in average between two and 10 blood feeding attempts during a day, i.e. between 0.2 and 0.8 hour^−1^ during the biting activity period, leading to an average of 0.5 hour^−1^. Consequently, the average biting rate by stable flies was assumed to be distributed according to a Pert distribution of parameters 0.4, 0.5 and 0.6 between 8am and 8pm and to be 0 otherwise.

Parametrizing *b_2_* (the probability that a vector becomes contaminated if it bites an infectious pig) requires parametrizing several parameters, including the viremia, the infectious dose for pigs and the crop volume in stable flies. Viremia in pigs infected by the Georgia 2007/01 ASFV strain can range between 10^3^ and 10^7^ 50% hemadsorbing doses (HAD_50_) per 1 mL (Guinat et al., 2014) but can be as high as 10^8^ HAD_50_/mL towards the end of their clinical evolution (Gallardo et al., 2017). Therefore, we assumed an average viremia of 10^5^ HAD_50_/mL. Under experimental conditions, a dose of 100 HAD_50_ injected intra-dermally led to the infection of pigs with a probability of 1 (Bernard et al., 2016). However, it was shown that doses of two to 10 viral particles were sufficient to infect a pig with other less virulent strains (Pan and Hess, 1984). To parametrize the model, we therefore assumed an infectious dose of 10 HAD_50_ and included this parameter in a sensitivity analysis. Furthermore, it was shown that the volume of blood stored in the crop after a blood meal could vary between 0.6 and 1.1μL (Lee and Davies, 1979). Consequently, with a viremia in pigs of 100 HAD_50_/μL and a volume of blood stored in the crop of 0.6 μL, the probability that at least 10 HAD_50_ (the assumed infectious dose for pigs) are swallowed and stored in the crop (*b_2_*) was assumed to be 1. Also, the viral concentration in the crop was assumed to be equal to the viremia (100 HAD_50_/μL).

Similarly, parametrizing *b_1_* (the probability that a pig becomes infected if bitten by an infective vector) required the additional parametrization of the regurgitation probability (hereafter referred to as *r*) and the regurgitated blood volume (*vol.r*). Under experimental conditions, it was observed that stable flies could regurgitate some blood stored in their crop, generating opportunities for pathogen transmission (Butler et al., 1977; Lee and Davies, 1979; Coronado et al., 2004). However, it is not clear if this regurgitation process occurs routinely at each blood meal, or if it is occasional and linked to the meal nature (blood versus sugar), as suggested by (Lee and Davies, 1979). To the best of our knowledge, neither *r* nor *vol.r* have been properly quantified and their descriptions found in the literature are scarce and have not been repeated in the past 40 years. (Kloft, 1992), as cited by (Hornok et al., 2020), suggested that the volume of blood regurgitated by stable flies does not exceed 180nL. We therefore assumed a value of 0.05 for *r* and 40 nL for *vol.r*, and included these two parameters in the sensitivity analysis. Assuming that the number of viral particles in the regurgitated blood volume is distributed according to a binomial distribution of parameters *vol.r* and the viremia, the probability *v* that at least 10 viral particles are injected to the host during a regurgitation was calculated at 8.10^−3^. Consequently, *b_1_* being the product of *v* and *r*, *b_1_* was given the value 4.10^−4^. Note that, we assumed that the volume of residual blood on the mouth parts was negligible when compared to the volume of blood potentially regurgitated from the crop, and so did not consider it in the model.

Assuming that ingested blood is stored in the crop for around 24h (Coronado et al., 2004; Baldacchino et al., 2013), the average duration of the infective period in stable flies (*τ*) was assumed to be 24h.

### Simulations

The model was initialised by introducing one exposed pig and simulated for 100 days in a deterministic framework. For a given scenario, 1000 simulations were run with parameters being randomly sampled in their respective distributions (Table 1) before each simulation. All scenarii combining pig-to-pig transmission and vector-borne transmission were compared to a scenario of pig-to-pig transmission only. The comparison was made on the duration between virus introduction and either (i) the moment when 10% of pigs were expected to be dead or (ii) the epidemic peak (i.e. the moment with the highest number of infectious pigs). Finally, for each scenario, the relative contribution of vector-borne transmission was calculated as the proportion of infection events that were due to the mechanical transmission by stable flies.

### Sensitivity analysis

A sensitivity analysis was conducted to investigate how the overall relative contribution of vector-borne transmission was influenced by the three input parameters associated with the most significant uncertainty: the infectious dose for pigs (values assessed: 1, 5, 10, 20 and 50 HAD_50_), the regurgitated blood volume (values assessed: 10, 20, 40, 100 and 180 nL) and the regurgitation probability (values assessed: 0.01, 0.05, 0.1 and 0.2). To do so, the value of each of these three parameters was changed individually while the value of the others was kept at their baseline values (Table 1), and 1000 model simulations were run.

## RESULTS

As depicted in Figure 2, assuming only pig-to-pig transmission, the time to reach 10% pig mortality would be around 32 days (interquartile range (IQR): 29 – 37) while the time to reach the epidemic peak would be around 49 days (IQR: 37 – 110).

**Figure 2:**
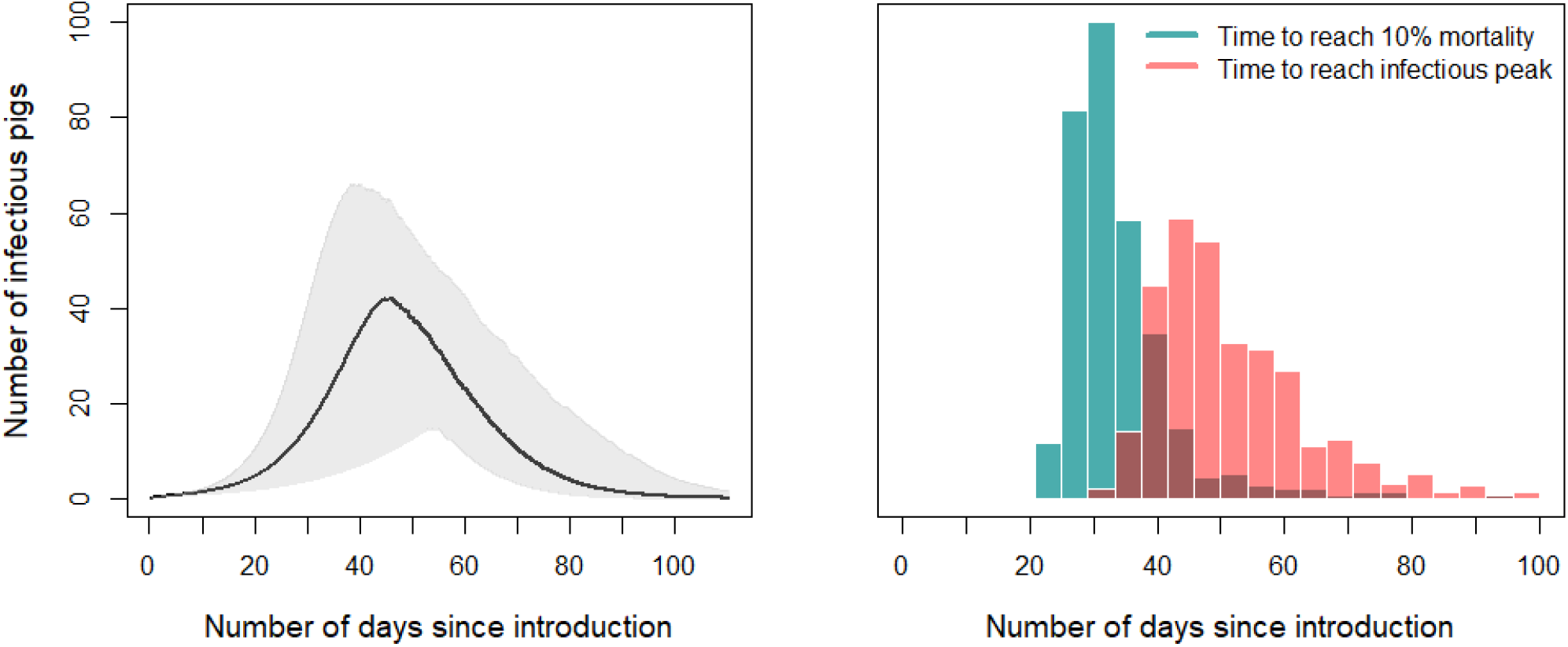
Expected within-farm African swine fever virus dynamic in the absence of stable flies. Left: Number of infectious pigs over time; the solid line corresponds to the median and the grey area to the 95% confidence region. Right: Distribution of the expected time to reach 10% mortality (green) and the infectious peak (red).

With the model assumptions, results suggest that in a context of a low density of stable flies in farms (<5 per pig), flies would play a minor role in the spread of ASFV as they are expected to be responsible for around 10% (IQR: 8 – 12%) of transmission events (Table 2). In this situation, the time to reach a pig mortality of 10% and the time to reach the epidemic peak would not be altered substantially as they would likely be reduced by only 10% (IQR: 8 – 13%) and 11% (IQR: 9 – 14%), respectively.

**Table 2:** African swine fever virus transmission model outputs for increasing abundance of stable flies.

| Number of stable flies per pig | Relative reduction of the time to reach a mortality of 10% (inter-quartile range) | Relative reduction of the time to reach the infectious peak (IQR) | Relative contribution of mechanical transmission by stable flies (IQR) |
| --- | --- | --- | --- |
| 5 | 0.10 (0.08 – 0.13) | 0.11 (0.09 – 0.14) | 0.10 (0.08 – 0.12) |
| 10 | 0.17 (0.13 – 0.22) | 0.19 (0.15 – 0.25) | 0.18 (0.15 – 0.21) |
| 20 | 0.26 (0.22 – 0.32) | 0.29 (0.24 – 0.35) | 0.29 (0.25 – 0.33) |
| 50 | 0.40 (0.36 – 0.47) | 0.44 (0.38 – 0.51) | 0.48 (0.43 – 0.52) |
| 100 | 0.52 (0.57 – 0.57) | 0.56 (0.50 – 0.62) | 0.64 (0.59 – 0.67) |

However, for an increased abundance of stable flies, model outputs indicate that their contribution could become substantial. Indeed, with densities of 20 and 50 flies per pig, the vector-borne transmission would likely be responsible for around 29% (IQR: 25 – 33%) and 48% (IQR: 43 – 52%) of transmission events, respectively (Table 2). In these two situations, the time to reach a pig mortality of 10% would be reduced by around 26% (IQR: 22 – 32%) and 40% (IQR: 36 – 47%), respectively (Figure 3).

**Figure 3:**
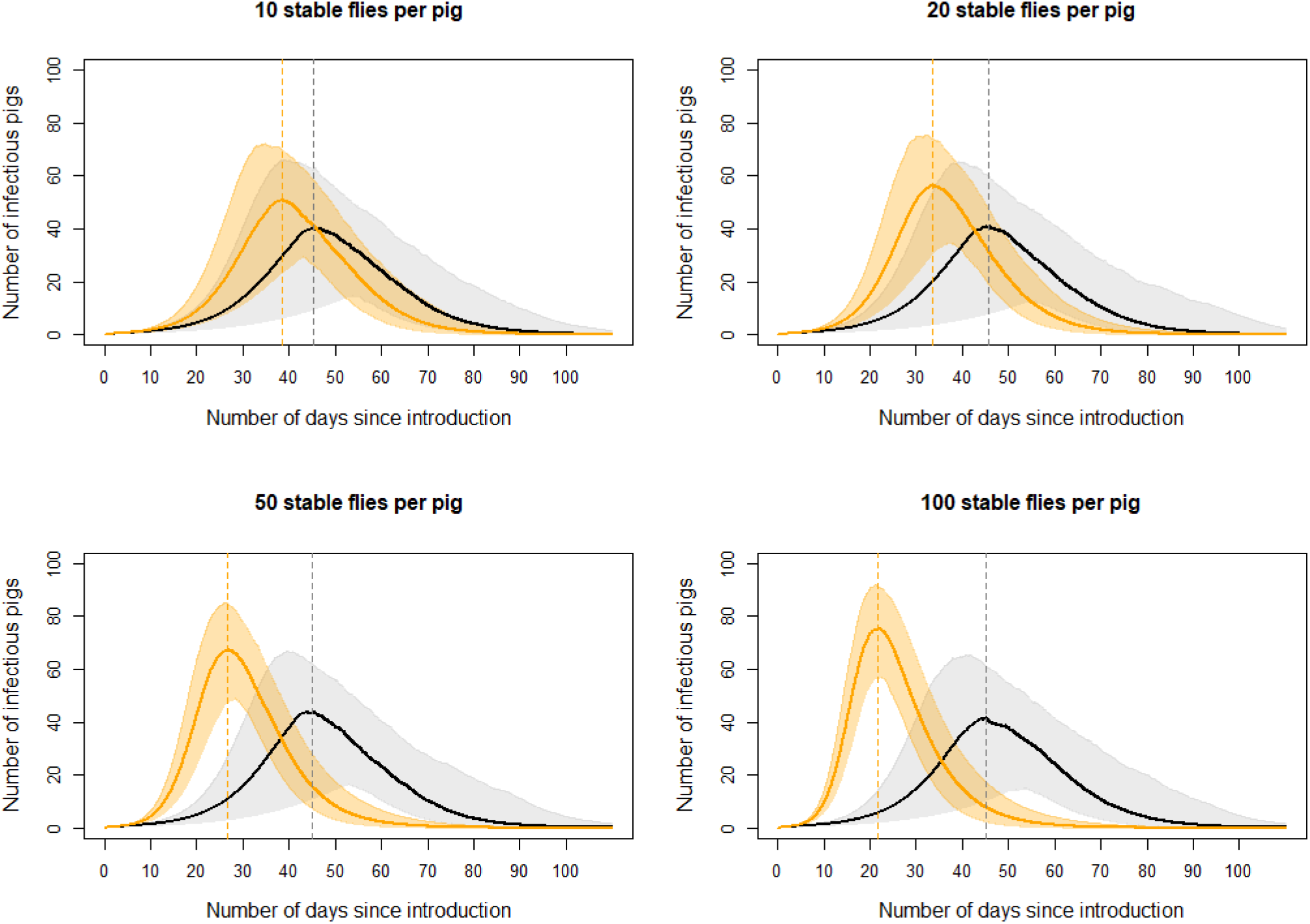
African swine fever virus transmission dynamics for increasing abundance of stable flies. Each panel corresponds to a different scenario of fly abundance from 10 per pig in the top left panel to 100 per pig in the bottom right panel); the orange part corresponds to a situation with the stable flies while the grey part corresponds to a situation without any stable fly; the solid lines correspond to the medians and the shadowed area to the 95% confidence regions.

The sensitivity analysis emphasised that the relative contribution of stable flies in virus transmission was strongly dependent on the assumptions made on the volume of regurgitated blood and on the infectious dose for pigs, irrespective of the number of stable flies per pig (Figure 4): when the volume of regurgitated blood or the infectious dose departed from their baseline assumption (40 nL and 10 HAD_50_, respectively), the model predicted that *Stable* flies would likely be responsible for almost no transmission (for a volume of regurgitated blood less than 20 nL or an infectious dose greater than 20 HAD_50_) or more than 95% of transmission events (for a volume of regurgitated blood greater than 100 nL or an infective dose less than 5 HAD_50_). The model appeared relatively less sensitive to the assumption made on the regurgitation probability (Figure 4).

**Figure 4:**
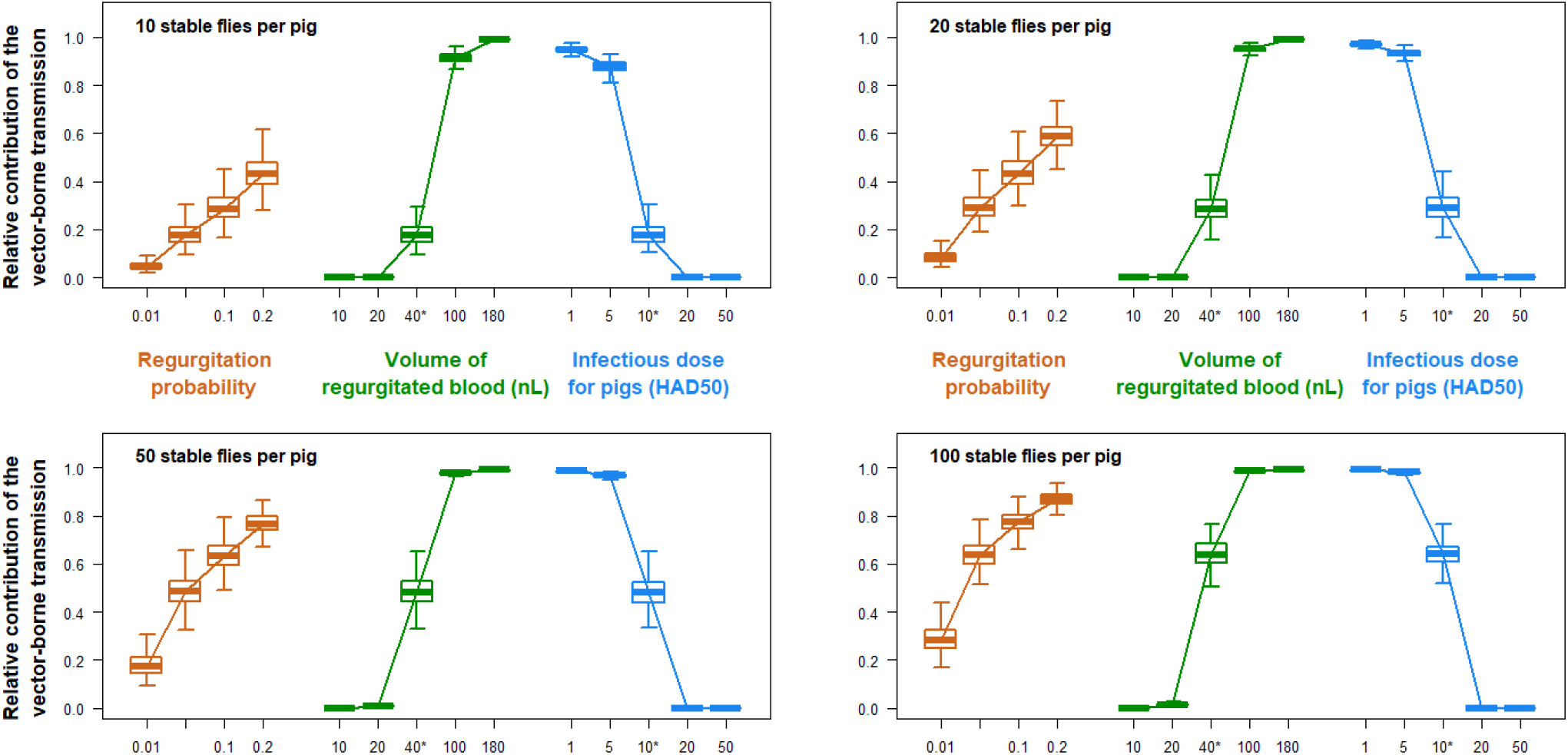
Impact of the regurgitation probability (brown), the volume of the regurgitated blood (green) and the infectious dose on the relative contribution of the stable flies-borne transmission of African swine fever virus (blue) for different densities of stable flies per pig. The wildcards denote the values assumed in the baseline model.

## DISCUSSION

Recent epidemiological investigations have raised serious questions regarding the role of blood-feeding arthropods in the spread of ASFV (Miteva et al., 2020). Experimental evidence has suggested that stable flies could play a role in its transmission (Mellor et al., 1987; Olesen, Hansen et al., 2018; Olesen, Lohse et al., 2018) and a recent prioritization exercise ranked them as the most probable blood-feeding arthropod to be a vector of ASFV in Metropolitan France (Saegerman et al., 2020). To gain insights into their potential contribution to the spread of the virus within a pig farm, we developed a mechanistic vector-borne transmission model of ASFV, for various densities of stable flies. Model outputs suggested that their contribution would be limited for low abundance of flies, but could increase dramatically with increasing fly abundance, substantially reducing the delay to reach the infectious peak in the farm. However, the model outputs being quite sensitive to some input model parameters (volume of regurgitated blood and infectious dose in pigs), experimental research is urgently required to better characterise their role.

Understanding the role of stable flies in ASFV transmission within an outdoor domestic pig farm is crucial for designing effective preventive and intervention strategies that could help decrease the risk of diffusion of the virus outside the farm. Indeed, if the time to reach a pig mortality of 10% was reduced by almost 20%, as the model suggested for a density of 10 flies per pig, farms with stable flies would exert a much stronger infectious pressure to other farms or to the neighbouring wildlife than farms free of stable flies. This would have three main consequences. First, it would imply that preventing the multiplication of stable flies in outdoor domestic pig farms would likely maintain a limited transmission rate of the virus, buying time to implement appropriate intervention measures in the infected premises. Secondly, reporting suspicions and planning interventions in a timely manner would be crucial in farms where stomoxes are present, as the model suggests that the increase in the number of infectious pigs would occur more rapidly than in farms without stable flies. Consequently, ensuring a high level of ASF awareness amongst pig owners whose farms present high densities of stable flies would be paramount. Finally, immediate implementation of vector control measures in infected premises could be necessary to curb the transmission rate in infected premises and decrease the risk of ASFV spread outside the farm.

The mechanistic model that was developed in this study allowed us to identify crucial knowledge gaps that need to be filled in order to assess precisely the potential contribution of stable flies to the spread of ASFV into the field. More evidence on the vectorial competence and capacity of stable flies to transmit ASFV mechanically is effectively paramount. First, it is necessary to reproduce experimental transmission assays on pigs using stable flies previously fed with infectious blood in order to confirm and analyse mechanical transmission of the virus. For example, the sensitivity analysis of the model demonstrated that it is essential to better understand blood-feeding behaviours of stable flies by quantifying both the frequency of regurgitation and the regurgitated blood volume. It would be also essential to run isolation tests on the blood stored in the crop of stable flies captured in the vicinity of infected farms. All of that would enable the quantification of the overall probability that an infective stable fly (i.e. partially fed with infectious blood) transmits the virus to a susceptible pig during a subsequent blood-feeding attempt (parameter *b_1_* in the current model). Also, it is necessary to describe quantitatively the populations of blood-feeding arthropods in pig farms and to identify farm characteristics that influence stable fly densities. Indeed, the literature related to the presence of stable flies in pig farms was very scarce, as we found only one publication on the subject (Moon et al., 1987).

Haematophagous arthropods biting infected wild boars – capable of roaming in close proximity to the farms – are also suspected to play a role in the introduction of ASFV into pig rearing facilities. The mechanistic model presented in this study could be extended to address this issue, though it would require several additional assumptions regarding ecological behaviour of wild boar with respect to pig facilities, ASFV prevalence within wild boar populations, and flying pattern of stable flies around pig facilities. Given the substantial amount of uncertainty already present on some parameters of the current model, we strongly believe that more precise knowledge on these parameters is required before such a mechanistic model could be useful to investigate the potential introduction of the virus into pig farms via stable flies.

## CONCLUSION

Although uncertainties remain on vector transmission of ASFV by blood-feeding arthropods, our model clearly highlights the threat these arthropods could pose by promoting the spread of the virus within outdoor pig farms. Further research is needed to address their role in the introduction of the virus into domestic pig facilities from infected wild boar populations. Such vector transmission would represent an additional indirect infection route that has barely been considered previously, generating additional control options that could complement standard regulated procedures such as culling, cleaning and disinfection.

## ACKNOWLEDGEMENTS

This assessment was conducted by the ad hoc working group “ASF vectors” of the French Agency for Food, Environmental and Occupational Health & Safety (Anses).

## ETHICAL APPROVAL

Due to the nature of the study and the low risk posed to participants, formal approval from an Ethics Committee was not a requirement at the time of the study.

## DATA SHARING AND ACCESSIBILITY

The data and codes that support the findings of this study are available from the corresponding author upon reasonable request.

## REFERENCES

Andraud, M., T. Halasa, A. Boklund, and N. Rose, 2019: Threat to the French Swine Industry of African Swine Fever: Surveillance, Spread, and Control Perspectives. Front Vet Sci 6, DOI: 10.3389/fvets.2019.00248.

Baldacchino, F., V. Muenworn, M. Desquesnes, F. Desoli, T. Charoenviriyaphap, and G. Duvallet, 2013: Transmission of pathogens by Stomoxys flies (Diptera, Muscidae): a review. Parasite 20, DOI: 10.1051/parasite/2013026.

Beltran-Alcrudo, D., J.R. Falco, E. Raizman, and K. Dietze, 2019: Transboundary spread of pig diseases: the role of international trade and travel. BMC Vet Res 15, DOI: 10.1186/s12917-019-1800-5.

Bernard, J., E. Hutet, F. Paboeuf, T. Randriamparany, P. Holzmuller, R. Lancelot, V. Rodrigues, L. Vial, and M.-F. Le Potier, 2016: Effect of O. porcinus Tick Salivary Gland Extract on the African Swine Fever Virus Infection in Domestic Pig. PLoS ONE 11, e0147869, DOI: 10.1371/journal.pone.0147869.

Berry, I.L., and J.B. Campbell, 1985: Time and Weather Effects on Daily Feeding Patterns of Stable Flies (Diptera: Muscidae). Environ Entomol 14, 336–342, DOI: 10.1093/ee/14.3.336.

Butler, J.F., W.J. Kloft, L.A. DuBose, and E.S. Kloft, 1977: Recontamination of food after feeding a 32P food source to biting muscidae. J. Med. Entomol. 13, 567–571, DOI: 10.1093/jmedent/13.4-5.567.

Coronado, A., J. Butler, J. Becnel, and J. Hogsette, 2004: Artificial feeding in the stable fly Stomoxys calcitrans and their relationship with the blood meal destination.. Presented at the 1st international symposium on Haemoparasites and their vectors, Caracas, Venezuela.

Cwynar, P., J. Stojkov, and K. Wlazlak, 2019: African Swine Fever Status in Europe. Viruses 11, 310, DOI: 10.3390/v11040310.

Fila, M., and G. Woźniakowski, 2020: African Swine Fever Virus - The Possible Role of Flies and Other Insects in Virus Transmission. J Vet Res 64, 1–7, DOI: 10.2478/jvetres-2020-0001.

Fischer, O., L. Mátlová, L. Dvorská, P. Svástová, J. Bartl, I. Melichárek, R.T. Weston, and I. Pavlík, 2001: Diptera as vectors of mycobacterial infections in cattle and pigs. Med. Vet. Entomol. 15, 208–211, DOI: 10.1046/j.1365-2915.2001.00292.x.

Gallardo, C., A. Soler, R. Nieto, C. Cano, V. Pelayo, M.A. Sánchez, G. Pridotkas, J. Fernandez-Pinero, V. Briones, and M. Arias, 2017: Experimental Infection of Domestic Pigs with African Swine Fever Virus Lithuania 2014 Genotype II Field Isolate. Transbound Emerg Dis 64, 300–304, DOI: 10.1111/tbed.12346.

Gilles, J., 2005: Dynamique et génétique des populations d’insectes vecteurs: les stomoxes, Stomoxys calcitrans et Stomoxys niger niger dans les élevages bovins réunionnais.

Guinat, C., A. Gogin, S. Blome, G. Keil, R. Pollin, D.U. Pfeiffer, and L. Dixon, 2016: Transmission routes of African swine fever virus to domestic pigs: current knowledge and future research directions. Vet Rec 178, 262–267, DOI: 10.1136/vr.103593.

Guinat, C., S. Gubbins, T. Vergne, J.L. Gonzales, L. Dixon, and D.U. Pfeiffer, 2016: Experimental pig-to-pig transmission dynamics for African swine fever virus, Georgia 2007/1 strain. Epidemiol Infect 144, 25–34, DOI: 10.1017/S0950268815000862.

Guinat, C., A.L. Reis, C.L. Netherton, L. Goatley, D.U. Pfeiffer, and L. Dixon, 2014: Dynamics of African swine fever virus shedding and excretion in domestic pigs infected by intramuscular inoculation and contact transmission. Vet Res 45, DOI: 10.1186/s13567-014-0093-8.

Halasa, T., A. Bøtner, S. Mortensen, H. Christensen, N. Toft, and A. Boklund, 2016: Simulating the epidemiological and economic effects of an African swine fever epidemic in industrialized swine populations. Vet. Microbiol. 193, 7–16, DOI: 10.1016/j.vetmic.2016.08.004.

Herm, R., L. Tummeleht, M. Jürison, A. Vilem, and A. Viltrop, 2019: Trace amounts of African swine fever virus DNA detected in insects collected from an infected pig farm in Estonia. Vet Med Sci 6, 100–104, DOI: 10.1002/vms3.200.

Hornok, S., N. Takács, S. Szekeres, K. Szőke, J. Kontschán, G. Horváth, and L. Sugár, 2020: DNA of Theileria orientalis, T. equi and T. capreoli in stable flies (Stomoxys calcitrans). Parasites & Vectors 13, 186, DOI: 10.1186/s13071-020-04041-1.

Kloft, W.J., 1992: Radioisotopes in vector researchSpringer., Vol. 9, pp. 41–66. In: Advances in disease vector research. New York.

Kukielka, E.A., B. Martínez-López, and D. Beltrán-Alcrudo, 2017: Modeling the live-pig trade network in Georgia: Implications for disease prevention and control. PLoS One 12, DOI: 10.1371/journal.pone.0178904.

Kunz, S.E., and J. Monty, 1976: Biology and ecology of Stomoxys nigra Macquart and Stomoxys calcitrans (L.) (Diptera, Muscidae) in Mauritius. Bulletin of Entomological Research 66, 745–755, DOI: 10.1017/S0007485300010798.

Lee, R.M.K.W., and D.M. Davies, 1979: Feeding in the Stable Fly, Stomoxys Calcitrans (Diptera: Muscidae) I. Destination of Blood, Sucrose Solution and Water in the Alimentary Canal, The Effects of Age on Feeding, And Blood Digestion. J Med Entomol 15, 541–554, DOI: 10.1093/jmedent/15.5-6.541.

Lichoti, J.K., J. Davies, P.M. Kitala, S.M. Githigia, E. Okoth, Y. Maru, S.A. Bukachi, and R.P. Bishop, 2016: Social network analysis provides insights into African swine fever epidemiology. Preventive Veterinary Medicine 126, 1–10, DOI: 10.1016/j.prevetmed.2016.01.019.

Mellor, P.S., R.P. Kitching, and P.J. Wilkinson, 1987: Mechanical transmission of capripox virus and African swine fever virus by Stomoxys calcitrans. Research in Veterinary Science 43, 109–112, DOI: 10.1016/S0034-5288(18)30753-7.

Miteva, A., A. Papanikolaou, A. Gogin, A. Boklund, A. Bøtner, A. Linden, A. Viltrop, C.G. Schmidt, C. Ivanciu, D. Desmecht, D. Korytarova, E. Olsevskis, G. Helyes, G. Wozniakowski, H.-H. Thulke, H. Roberts, J.C. Abrahantes, K. Ståhl, K. Depner, L.C.G. Villeta, M. Spiridon, S. Ostojic, S. More, T.C. Vasile, V. Grigaliuniene, V. Guberti, and R. Wallo, 2020: Epidemiological analyses of African swine fever in the European Union (November 2018 to October 2019). EFSA Journal 18, e05996, DOI: 10.2903/j.efsa.2020.5996.

Moon, R.D., L.D. Jacobson, and S.G. Cornelius, 1987: Stable flies (Diptera: Muscidae) and productivity of confined nursery pigs. J. Econ. Entomol. 80, 1025–1027, DOI: 10.1093/jee/80.5.1025.

Olesen, A.S., M.F. Hansen, T.B. Rasmussen, G.J. Belsham, R. Bødker, and A. Bøtner, 2018: Survival and localization of African swine fever virus in stable flies (Stomoxys calcitrans) after feeding on viremic blood using a membrane feeder. Vet. Microbiol. 222, 25–29, DOI: 10.1016/j.vetmic.2018.06.010.

Olesen, A.S., L. Lohse, M.F. Hansen, A. Boklund, T. Halasa, G.J. Belsham, T.B. Rasmussen, A. Bøtner, and R. Bødker, 2018: Infection of pigs with African swine fever virus via ingestion of stable flies (Stomoxys calcitrans). Transbound Emerg Dis 65, 1152–1157, DOI: 10.1111/tbed.12918.

Oliveira, R.P. de, E. Hutet, F. Paboeuf, M. Duhayon, F. Boinas, A.P. de Leon, S. Filatov, L. Vial, and M.-F.L. Potier, 2019: Comparative vector competence of the Afrotropical soft tick Ornithodoros moubata and Palearctic species, O. erraticus and O. verrucosus, for African swine fever virus strains circulating in Eurasia. PLOS ONE 14, e0225657, DOI: 10.1371/journal.pone.0225657.

Pan, I.C., and W.R. Hess, 1984: Virulence in African swine fever: its measurement and implications. Am. J. Vet. Res. 45, 361–366.

Penrith, M.-L., A.D. Bastos, E.M.C. Etter, and D. Beltrán-Alcrudo, 2019: Epidemiology of African swine fever in Africa today: Sylvatic cycle versus socio-economic imperatives. Transbound Emerg Dis 66, 672–686, DOI: 10.1111/tbed.13117.

Pérez-Sánchez, R., A. Astigarraga, A. Oleaga-Pérez, and A. Encinas-Grandes, 1994: Relationship between the persistence of African swine fever and the distribution of Ornithodoros erraticus in the province of Salamanca, Spain. Vet. Rec. 135, 207–209, DOI: 10.1136/vr.135.9.207.

Petrasiunas, A., R. Bernotiene, and J. Turcinaviciene, 2018: Catches of blood-feeding flies with NZI traps in African swine fever affected areas of Lithuania. Bulletin of the Lithuanian entomological society.

Podgórski, T., and K. Śmietanka, 2018: Do wild boar movements drive the spread of African Swine Fever? Transbound Emerg Dis 65, 1588–1596, DOI: 10.1111/tbed.12910.

Quembo, C.J., F. Jori, L. Heath, R. Pérez-Sánchez, and W. Vosloo, 2016: Investigation into the Epidemiology of African Swine Fever Virus at the Wildlife - Domestic Interface of the Gorongosa National Park, Central Mozambique. Transbound Emerg Dis 63, 443–451, DOI: 10.1111/tbed.12289.

Quembo, C.J., F. Jori, W. Vosloo, and L. Heath, 2018: Genetic characterization of African swine fever virus isolates from soft ticks at the wildlife/domestic interface in Mozambique and identification of a novel genotype. Transboundary and Emerging Diseases 65, 420–431, DOI: 10.1111/tbed.12700.

Saegerman, C., S. Bonnet, E. Bouhsira, N. De Regge, J. Fite, F. Etore, M.-M. Garigliany, F. Jori, L. Lempereur, M.-F. Le Potier, E. Quillery, T. Vergne, and L. Vial, 2020: An expert opinion assessment of blood-feeding arthropods based on their capacity to transmit African swine fever virus in Metropolitan France. Submitted.

Salem, A., M. Franc, P. Jacquiet, E. Bouhsira, and E. Liénard, 2012: Feeding and breeding aspects of Stomoxys calcitrans (Diptera: Muscidae) under laboratory conditions. Parasite 19, 309–317, DOI: 10.1051/parasite/2012194309.

Sánchez-Cordón, P.J., M. Montoya, A.L. Reis, and L.K. Dixon, 2018: African swine fever: A re-emerging viral disease threatening the global pig industry. Vet. J. 233, 41–48, DOI: 10.1016/j.tvjl.2017.12.025.

Sumner, T., R.J. Orton, D.M. Green, R.R. Kao, and S. Gubbins, 2017: Quantifying the roles of host movement and vector dispersal in the transmission of vector-borne diseases of livestock. PLoS Comput. Biol. 13, e1005470, DOI: 10.1371/journal.pcbi.1005470.

Turner, J., R.G. Bowers, and M. Baylis, 2012: Modelling bluetongue virus transmission between farms using animal and vector movements. Sci Rep 2, DOI: 10.1038/srep00319.

Vial, L., E. Ducheyne, S. Filatov, A. Gerilovych, D.S. McVey, I. Sindryakova, S. Morgunov, A.A. Pérez de León, D. Kolbasov, and E.M. De Clercq, 2018: Spatial multi-criteria decision analysis for modelling suitable habitats of Ornithodoros soft ticks in the Western Palearctic region. Vet. Parasitol. 249, 2–16, DOI: 10.1016/j.vetpar.2017.10.022.

